# The role of family and environment in determining the skin microbiome of captive aquatic frogs, *Xenopus laevis*

**DOI:** 10.1101/2023.10.05.561135

**Authors:** Phoebe A. Chapman, Daniel Hudson, Xochitl C. Morgan, Caroline W. Beck

**Affiliations:** Department of Zoology, University of Otago, Dunedin, New Zealand; Department of Microbiology and Immunology, University of Otago, Dunedin, New Zealand; Department of Biostatistics, Harvard T. H. Chan School of Public Health, Boston, Massachusetts, USA

**Keywords:** *Xenopus laevis*, microbiome, bacterial colonisation, 16S rRNA, Illumina MiSeq, amphibians

## Abstract

**Background:** The amphibian skin microbiome has drawn interest due to the ecological threat posed by chytridiomycosis, which drives changes in symbiotic microbial communities and may be inhibited by certain bacterial taxa. However, skin microbes also play a role in amphibian tissue regeneration. *Xenopus* spp. are well-established model organisms used to study development, regeneration, genetics and disease. Husbandry protocols, including use of antibiotics and other sterilising agents, may affect experimental outcomes by altering microbiomes. It is therefore essential to improve our understanding of *Xenopus* microbiome characteristics and inheritance. We undertook bacterial 16S rRNA based sampling of a captive, closed *Xenopus laevis* colony. A total of 16 female frogs, their eggs, and tadpoles were sampled, covering multiple aquarium systems and tanks, along with testes from males used for in vitro fertilisation and a range of environmental samples.

**Results:** Tank environments supported the most complex microbial communities. Mother frogs harboured the most diverse microbial communities of the frog life stages, with tadpole skin microbiomes being relatively simple. Frog samples were dominated by Proteobacteria and Bacteroidota. *Rhizobium* and *Chryseobacterium* were dominant in tadpoles, whereas mothers supported high proportions of *Vogesella* and *Acinetobacter* as well as *Chryseobacterium*. While the mothers’ habitats contained low levels of these taxa, the tadpole’s environmental microbes were very similar to those on tadpole skin. A total of 34 genera were found to be differentially abundant between the mothers and tadpoles. Analysis of Bray-Curtis distances indicated that mother and tadpole microbiomes varied according to the mothers’ aquarium system, the tanks within them, and the individual mother. Source tracking analyses showed that egg jelly and tadpoles received a mean of approximately two thirds of their microbiomes via vertical transmission, although a sizeable proportion came from unknown sources at all life stages.

**Conclusions:** The skin of mother frogs appears to select for certain taxa that are otherwise present at low abundances in the environment. While tadpoles inherit a proportion of their microbiomes from their mothers via the egg, they support a distinct and less diverse microbial community than adult frogs. The microbiome varies between individual mothers, and is also affected by the aquarium system and individual tank within that the mother occupies.

## 1 Background

Symbiotic microbe communities are important to the function and health of their host. They instigate and mediate immune, metabolic, developmental and other physiological pathways in a broad range of host taxa, from plants [1, 2] through to invertebrates [3, 4] and vertebrates [5, 6], including humans [7, 8]. In amphibians, the skin is an organ of particular importance due to its roles in gas exchange, osmoregulation and immune defence [9, 10]. Microbes inhabiting the amphibian skin might influence their co-habiting community by producing antimicrobial compounds, stimulating the secretion of antimicrobial peptides from the host’s cutaneous mucous glands, or simply by competing for resources [11–13]. Due to conservation problems caused by the fungal pathogen *Batrachochytrium* spp., many amphibian microbiome studies have focused on the role of skin-dwelling bacterial communities in disease susceptibility (see for examples [14–17]). Some have attempted to determine how and from where microbes colonise the amphibian integument, however conclusions vary. Several studies have found that host species was the most important predictor of skin microbiome composition among aquatic species, with co-habitation having little effect [18, 19]. However, others [20–22] have found that the environment is a significant, if not the primary determinant of skin microbiome assemblages (reviewed by [23]). Differences in study design (e.g. use of high throughput sequencing vs. culture based or other methods) and taxa/taxonomic levels (salamanders and newts vs. frogs; species vs. order) may account for some of these contrasting conclusions. While the microbiome may be seeded from various sources, including the habitat and vertical or horizontal transfer from conspecifics, the amphibian skin potentially selects for and provides a competitive advantage for certain microbes that may otherwise be encountered in low numbers [19].

While the amphibian skin microbiome has been the subject of various studies in relation to chytridiomycosis, habitat and host taxonomy, its effects on physiological pathways have received considerably less attention. Bacterial communities play a critical role in activating pathways involved in development, limb regeneration and wound healing in model organisms from other taxonomic groups, including planaria [24, 25], mice [26, 27] and axolotls [28]. Clawed frogs from the *Xenopus* genus are proven and widely used models for laboratory developmental and regeneration studies. Recently, it has become evident that bacteria are important in *Xenopus* tail regeneration [29, 30]. While Mashoof et al. [31] were the first to characterize the *Xenopus laevis* gut microbiome, and demonstrated that thymectomy did not affect its composition, Scalvenzi et al. [32] found that the size and composition of *Xenopus tropicalis* gut microbiomes vary according to developmental stage. This can be at least partially explained by changes in diet and gut structure at different stages of ontogeny, however the potential role of these bacteria in influencing the developmental process is yet to be properly explored. Li et al. [33] demonstrated that the gut microbiome of *X. tropicalis* froglets was distinct from their food and surrounding water, suggesting that frogs at this life stage may exhibit a degree of immune control over their microbiome composition. Intriguingly, Bishop & Beck [29] found that exposure to various antibiotics, commonly used by *Xenopus* breeding facilities during the embryo rearing process, resulted in a sharp decrease in tail regeneration success in *Xenopus laevis* tadpoles. Meanwhile, the addition of either heat-killed *Escherichia coli* or commercially supplied lipopolysaccharide (LPS) from *E. coli* or *Pseudomonas aeruginosa* improved the likelihood of tail regeneration, including following antibiotic treatment. Further to this, Chapman et al. [30] characterized microbiomes from *X. laevis* tadpoles and demonstrated that LPS from other gram-negative bacteria, including commensal taxa, can promote regeneration, with some taxa over-represented in successful regenerators. Piccinni et al. [34] compared microbial communities living on ‘clean’ (thimerosal/ethanol treated) and standard feeding stage *X. laevis* tadpoles and adults, finding that microbiomes between these life stages were distinct. Tadpoles hosted a relatively diverse microbe assemblage which correlated with their environment (tank water), while those on adult frogs were less varied and showed little similarity to the surrounding environment, supporting the theory that amphibians acquire an ability to moderate their microbiome through immune or other means as they develop. Adult frog microbiomes were not significantly affected by housing conditions (‘clean’ vs standard), however some differences in beta diversity and microbe community composition were observed in tadpoles.

While Piccinni et al. [34] demonstrated the characteristics of *X. laevis* skin microbiomes at two key life stages, their tadpoles were not descended from the adults studied and were routinely raised in antibiotics, so inferences on maternal transfer of skin microbiota (i.e. vertical transfer) were not possible. Chapman et al. [30] compared microbiomes between treated and untreated tadpole clutches (sibships) from different mothers and noted considerable variation between sibships. The relative roles of vertical familial transmission and the environment on the microbiome at various stages of the lifecycle remain to be explored. As *Xenopus* spp. are widely used as models in regeneration and developmental studies, and intervention with their microbiomes influences regenerative outcomes [29, 30], it is critical to understand how their microbiota vary across individuals and colonies, and where they come from. To answer these questions, we characterised and compared the bacterial communities present on adults, eggs and tadpoles from non-antibiotic treated *X. laevis* using 16S ribosomal RNA amplicon sequencing. We also sequenced samples taken from potential environmental sources to determine their contribution at various *X. laevis* life stages.

## 2 Materials and Methods

### 2.1 *Xenopus* husbandry

The *X. laevis* colony at the University of Otago, New Zealand is comprised of several recirculating aquarium systems, each comprised of a number of smaller tanks and supplied by carbon filtered town tap water. The colony is closed, with no frogs introduced from outside sources since 2004, and current adults are captive-bred F_1_ or F_2_ generations. Frogs are fed with generic salmon farming pellets twice weekly. The aquarium system uses bioballs as biological filtration, which are plastic spheres with a network of channels and ridges designed to provide maximum surface area for nitrifying bacteria.

### 2.2 Embryo production and sample collection

All samples were collected in 2020. Eggs and embryos from sixteen female frogs were included in the study, with four females randomly selected from each of four different tanks split across two recirculating aquarium systems (**SUPP. FIGURE S1**). While frogs from a tank were not necessarily related, they were of similar age, with frogs from Tank B being considerably older (13 years) than those from Tanks C – E (2 - 4 years). Induction of egg laying, in vitro fertilisation, and embryo husbandry were as described by Chapman et al. [30]. Each female was placed in her own 1 L container of 1x MMR at 22 °C the morning following induction injection, allowing her eggs to be kept separate from those of others. Each tadpole sibship (family group derived from a particular mother) was split into two petri dishes containing 24 tadpoles each (total 48 tadpoles per sibship) with 30 ml 0.1x MMR (Marc’s Modified Ringer’s solution, pH 7.4: 100 mM NaCl, 2 mM KCl, 1 mM MgSO4.7H2O, 2 mM CaCl2, 5 mM HEPES, 0.1 mM EDTA at pH 8.0). Tadpole sibships did not have contact with other sibships at any stage, nor were they fed or in contact with any antibiotic or sterilising agent.

During the egg production stage above, batches of 10 freshly laid, unfertilised eggs were collected from each mother, and placed in a 1.5 ml Eppendorf tube (containing 200 µl filtered NaCl/Tween as below). Eggs were gently vortexed for 30 sec to dislodge bacteria on the jelly coats, and the sample pulsed in a centrifuge to enable retrieval of the supernatant. Tadpole samples were excised tail tips (one third of tail length) collected at developmental stage 46 [35] during a concurrent regeneration experiment. The removed tail portions were placed into 0.2 ml DNase/RNase free PCR strip tubes with 50 μl of filter-sterilised sodium chloride/Tween solution (0.15 M NaCl, 0.1% Tween20) and vortexed for 1 minute. A total of 6 tails were collected from each sibship. Unmoistened Puritan DNA-free cotton-tipped applicators were used to swab inner tank walls, the surfaces of bioballs, tadpole petri dishes, and adult female frogs. For the latter, each frog was swabbed on the dorsal and ventral surface, thigh crease and foot web with one swab, then repeated twice more with new swabs; each swab covered approximately one quarter of the animal’s surface area. Male frogs were sacrificed as part of the standard fertilisation protocol (refer Chapman et al. [30]), with one testis used to fertilise eggs from a female tank group while the second was divided into three pieces for microbial sequencing. Samples of tank water (50 ml), frog food (commercial salmon pellets), a bacterial aquarium conditioning additive (Stresszyme, API Fish Care) (200 µl), filtered Tween solution (200 µl), and tadpole raising media (0.1x MMR with residual 1/4000 v/v MS222 anaesthetic agent; 5 ml / sample), were also collected. Petri dish swabs and tadpole media were collected at the end of the regeneration experiment, one week after embryo fertilisation. All tank associated samples were collected at the time of female swabbing, with swabbing/experiments being completed on a tank-by-tank basis over the course of approximately two months. In all cases except tadpole tails, three replicates were collected (**SUPP. FIGURE S1**). Samples were stored at -20 °C until DNA extraction. A schematic of the summarised experimental design and sampling is provided as **FIGURE 1**.

**Figure 1:**
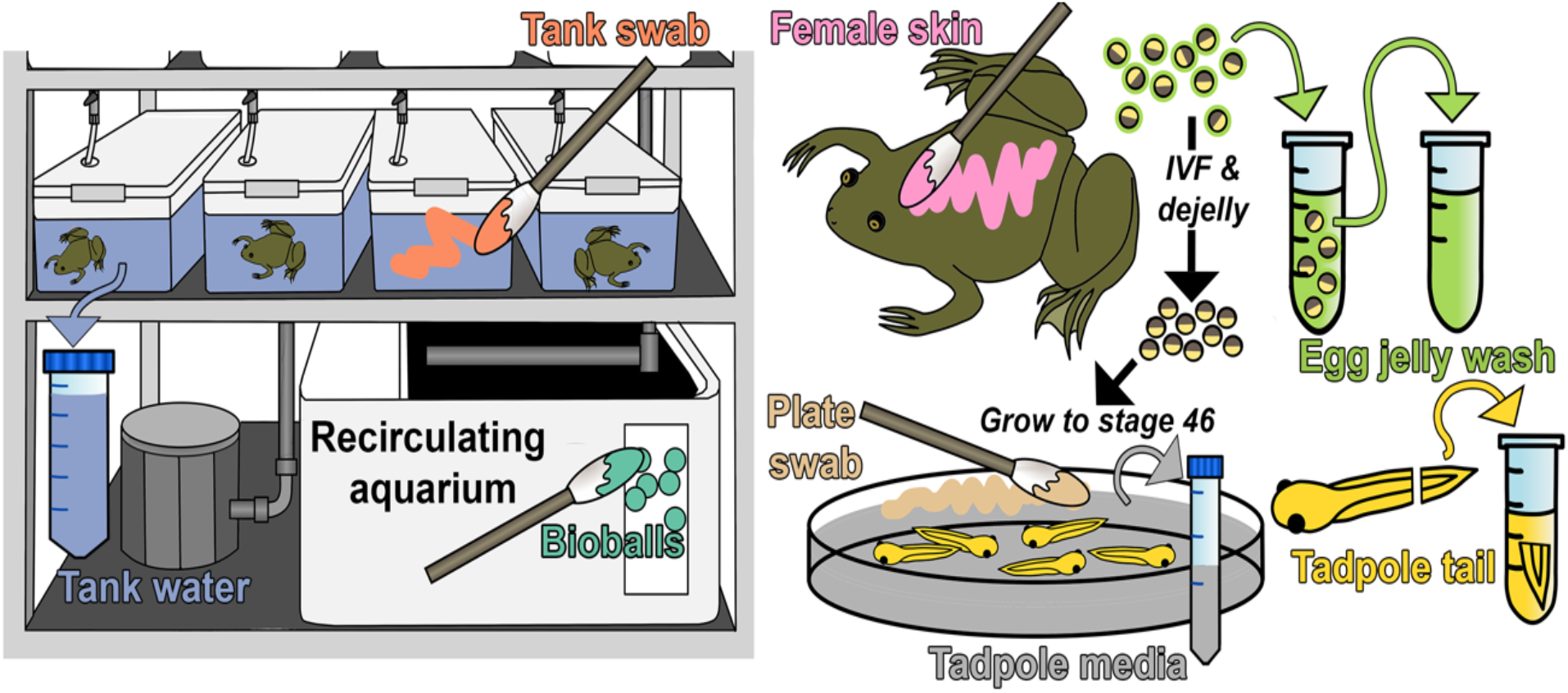
Summarised sampling strategy for study. For each tadpole sibship, 48 tadpoles were split between two petri dishes (plates), reared to stage 46 and sampled. Several samples that were shown to contribute negligable microbes (Stresszyme/frog food/testis) have been omitted from this diagram as preliminary findings suggested these do not contribute to tadpole microbial ecology. All samples were collected in triplicate and DNA extracted for amplification and sequencing of the 16S rRNA V4 hypervariable region. Further details are contained in **SUPP. FIGURE S1**.

### 2.3 DNA extraction and 16S rRNA amplicon sequencing

DNA was extracted from all samples described in **SECTION 2.2** along with the following controls: blank Puritan swab, blank filter (Millex GV 0.22 μm PVDF membrane – used to filter Tween prior to use), Tween, mock community (ZymoBIOMICS® Microbial Community Standard), and blank (no sample) extractions.

A DNeasy PowerLyser PowerSoil DNA extraction kit (Qiagen) was used to obtain DNA from samples as per manufacturer’s instructions. DNA was eluted into a final volume of 30 μl and stored at -80 °C. Amplification and sequencing of the 16S V4 hypervariable region was completed on the Illumina MiSeq platform by the Argonne National Laboratory (Illinois, USA). Sequencing was completed as per Caporaso et al. [36] using primers 515F/806R [37, 38] and peptide nucleic acid (PNA) PCR clamps to inhibit amplification of host mitochondrial sequences, resulting in paired end reads of 2 x 250 bp.

### 2.4 Sequence processing

Sequences were demultiplexed using idemp [39] and quality filtering and adaptor removal completed using fastp (v0.23.2) [40]. The DADA2 pipeline (v1.16) [41] was used to further filter and trim sequence data, with paired reads merged and chimeras removed before taxonomic assignment using the naïve Bayesian classifier method with reference to the SILVA database (v138.1). The dataset was then passed to R (v4.2.2) for further processing and subsequent analysis. Data was formatted into a phyloseq (v1.42.0) [42] object, and decontam (v1.18.0) [43] used to identify and remove potential contaminant amplicon sequence variants (ASVs) with reference to negative controls. ASVs were then agglomerated at genus level. A multiple sequence alignment was created using DECIPHER (v2.26.0) [44] and a maximum likelihood phylogenetic tree constructed with IQTree (v2.2.2.3) [45]. The TVMe+R9 substitution model was selected based on a best-fit assessment using the inbuilt ModelFinder [46] utility; 1066 bootstrap replicates were required for convergence.

Pairwise Pearson’s correlation coefficients were calculated for the three replicates within each sample, and the replicate with the lowest mean correlation coefficient removed from further analysis. The remaining two replicates were merged, with OTU counts averaged (speedyseq v0.5.3) [47]. The merged samples were then used for downstream analysis. For tadpole tails, the three individual tadpoles taken from each plate were treated as sample replicates.

After sequence processing and replicate merging, we identified 441 bacterial genera across 149 samples. Read counts in all control samples (excluding mock community samples) were negligible. Mock community profiles reflected those expected. A summary of read counts for each sample type is provided in **SUPP. TABLE S1**. Following an initial analysis of read counts and composition, samples for further analysis were narrowed down to a ‘core’ group of eight sample types comprised of bioballs, tank water, tank wall swabs, mother frog skin swab, egg jelly wash, tadpole tails, tadpole media and tadpole plate swabs.

### 2.5 Data Analyses

All analyses were completed using R v4.2.2, with statistics calculated using the base Rstats package unless otherwise specified.

#### 2.5.1 Alpha diversity

Rarefaction curves were prepared prior to analysis. Samples were subsequently rarefied to 1000 reads based on examination of rarefaction curves and read count distribution histograms. Observed, Shannon and Faith’s phylogenetic diversity were calculated for each sample type using phyloseq (Observed/Shannon) and picante v1.8.2 [48] (Faith’s). Kruskal-Wallis tests and post-hoc Dunn’s test with Bonferroni correction (the latter with FSA v0.9.4 [49]) were used to find significant differences between sample types following Shapiro-Wilk test of normality. Diversity metrics were visualised as boxplots created using ggplot2 (v3.4.0) [50].

#### 2.5.2 Beta diversity

Bray-Curtis distances were calculated between unrarefied samples using the phyloseq::distance function. Samples with less than 500 reads were excluded. Hopkins Test (clustertend v1.6) [51] and a silhouette analysis (factoextra v1.0.7) [51] were calculated for Bray-Curtis distances to assess the degree of clustering, and non-metric multidimensional scaling (NMDS) ordinations were prepared using phyloseq and ggplot2. Permutational multivariate analysis of variance (PERMANOVA – vegan v2.6-4 (adonis2 function) [52], and Wilcoxon Rank Sum test (rstatix v0.7.2) [53] or Kruskal-Wallis test were used to compare Bray-Curtis distances between sample types. Mother ID, tank, and aquarium system were designated as fixed effects with interactions for PERMANOVAs dependent on the comparison being made.

Heatmaps of Pearson’s correlation coefficients for Bray-Curtis distances between groups of samples were generated using corrplot v.0.92 [54].

#### 2.5.3 Community composition and differential abundance

Rarefied relative abundance of microbial taxa in sample types was visualised using the microshades (v1.0) [55] package. As a small number (6) of merged samples were lost in the process of rarefying, these samples were added back for the purpose of these plots only in order to enable comparisons between all mothers and their tadpoles. This was achieved by identifying the replicate with the highest read count and merging it with the most highly correlated of the remaining two replicates (based again on Pearson’s correlation coefficient) and discarding the third. In all but one case this was sufficient to enable rarefying to 1000 reads. This final sample was represented in relative abundance plots using unrarefied relative abundance values based on the two replicates with the highest read counts, which also exhibited the highest mean correlation coefficients.

The ANCOMBC2 (v2.0.1) [56] and MaAsLin2 (v1.12.0) [57] packages were used to calculate differential abundance of genera between key sample types, using unrarefied data with significance (alpha) set at 0.25. For ANCOMBC2, System, Tank, and Mother ID, and interactions between these predictors, were incorporated into the model, while a similar model was specified for MaAsLin2 without interactions. Microbial taxa found to be differentially abundant by both packages were extracted and mean relative abundance visualised across all sample types with ggplot2.

#### 2.5.4 Microbial source tracking

The FEAST package (v0.1.0) [58] was used to determine the likely origin of microbial taxa found in different sample types. Analyses were completed on unrarefied data in four broad groups (**TABLE 1**) selected based on likely direct interaction between samples. Where relevant, analyses were run individually for subgroups (for e.g. Tank B mothers with relevant Tank B environmental samples) and averaged for visualisation in a sankey plot using networkD3 (v0.4) [59]. After initial analysis we determined that tadpoles were most likely to be seeding their environment rather than vice versa as tadpoles shared a number of taxa with mother skin and egg jelly. Additionally, the MMR media, while not sterile after opening, was autoclaved prior to use.

**TABLE 1:** Groupings of sources and sinks for FEAST analyses.

| Group | Source/s | Sink |
| --- | --- | --- |
| 1 | Tank water<br>Tank swab | Mother skin swab |
| 2 | Mother skin swab | Egg jelly wash |
| 3 | Egg jelly wash | Tadpole tail |
| 4 | Tadpole tail<br>Egg jelly wash | Tadpole media |

## 3 Results

### 3.1 Different *Xenopus* life stages and environments support distinct microbial communities

Sample types, or groups of sample types, were broadly distinguishable based on their microbial communities. Samples associated with the adult frog tank environment (i.e. tank water, tank wall swabs, and bioballs) were the most complex, with tadpole environment samples (media and plate swabs) being least complex (**FIGURE 2A** and **SUPP. FIGURE S2A-B; SUPP. TABLES 2A-C)**. Frog samples (mother skin swabs, egg jelly, and tadpole tail clips) fell in between the environmental samples, with tadpole samples being consistently the least complex. This pattern remained generally consistent across all alpha diversity metrics calculated. Sample type accounted for 28% of variation in beta diversity based on Bray-Curtis distances (PERMANOVA (vegan::adonis2), p = 0.001, R^2^ = 0.276; **SUPP. TABLE S3**) while the aquarium system accounted for 2.8% (p = 0.001, R^2^ = 0.028) and the interaction between system and tank 8.6% (p = 0.001, R^2^ = 0.086). Hopkins statistic calculated on Bray-Curtis distances (0.161) indicated clustering within the data, with cluster (silhouette) analysis identifying two clusters (**SUPP. FIGURE S3**). NMDS ordination (**FIGURE 2B**) showed that aquarium surface samples (i.e. bioballs and tank swabs) overlapped substantially, forming a cluster distinct from tank water samples and the various *Xenopus* samples and tadpole environmental samples. Mother skin swabs and egg jelly samples clustered together, as did tadpole samples, tadpole plate swabs and media.

**Figure 2.**
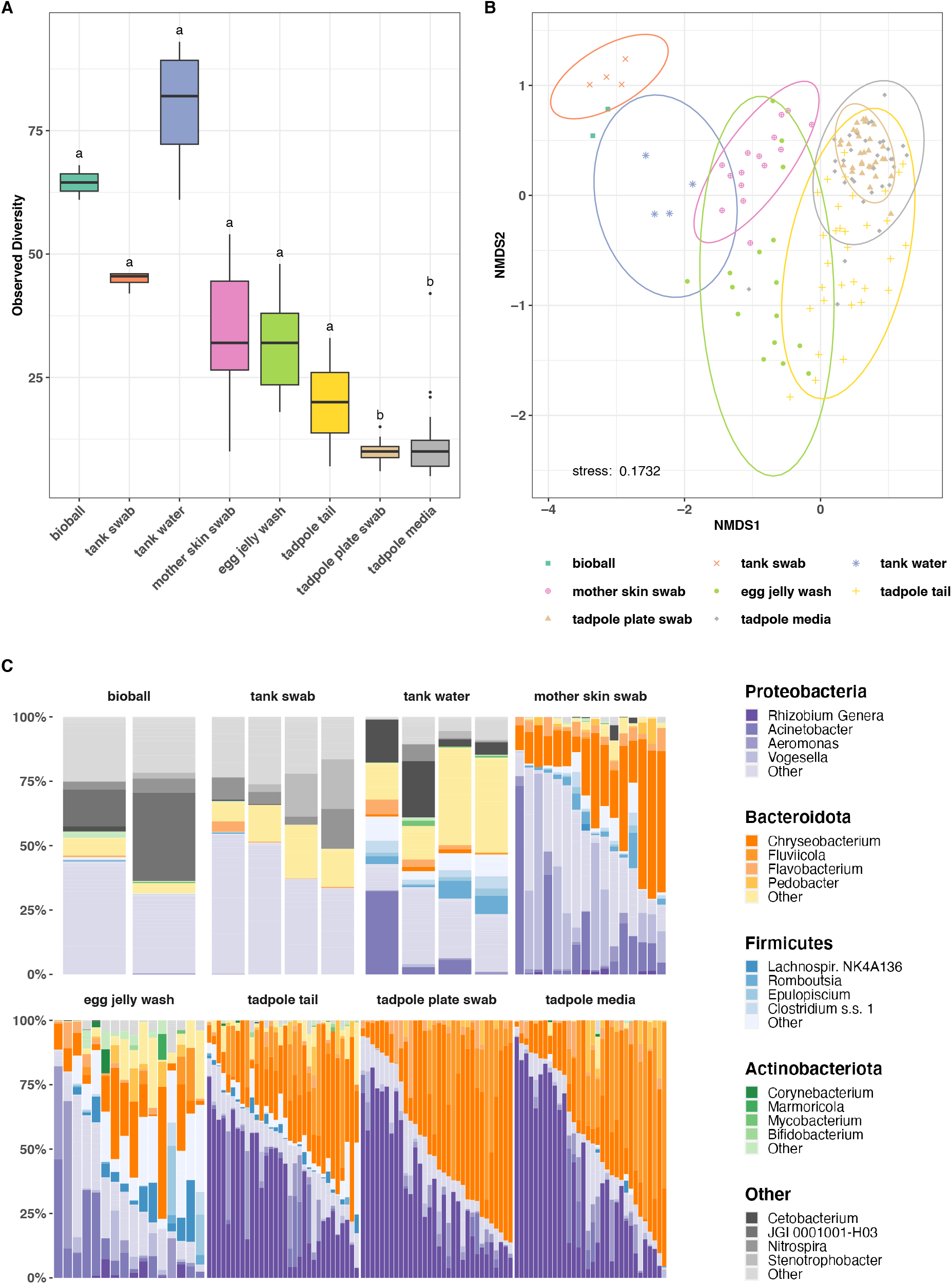
Comparison of microbial communities in core sample types based on 16S amplicon sequencing. A) Observed alpha diversity (rarefied at 1000 reads). Letters above each sample type denote statistical significance in compact letter display format; sample types with the same letter designation have no statistically significant difference (p < 0.05). B) Non-metric multidimensional scaling (NMDS) ordination of Bray-Curtis distance matrix of microbial communities from core sample types. C) Relative abundance of microbial phyla and genera in core sample types (rarefied at 1000 reads).

Overall, Proteobacteria and Bacteroidota were the most dominant bacterial phyla (**FIGURE 2C, TABLE 2**). When comparing frog associated samples, the majority of genera were present in at least two of the three samples types, with only 1, 1, and 2 of 87 total genera being unique to mother skin, egg jelly or tadpoles respectively. However, in most cases, abundances of genera differed markedly based on the sample type. *Chryseobacterium* (Bacteroidota) was common across all frog associated samples, being either the most or second most abundant taxa in mother skin, egg jelly and tadpole tail samples (**TABLE 2**), and was also common in the tadpole environment. *Vogesella* (Proteobacteria) was the second most abundant taxa on mother skin, while *Rhizobium* spp. (assigned as *Allorhizobium-Neorhizobium-Pararhizobium-Rhizobium* by the SILVA (v138.1) database, summarised here as *Rhizobium* genera for brevity) (Proteobacteria) was dominant in tadpole samples, followed by *Chryseobacterium* and *Fluviicola*. These latter three taxa also dominated tadpole environment samples. The adult frog environment shared relatively few taxa with tadpole tails or their associated environment, although *Acinetobacter* (Proteobacteria) was common in adult tank water, as well as in mother and egg samples. The top 5 taxa by relative abundance for frog samples are provided in **TABLE 2**. Top 5 genera for all core sample types are provided in **SUPP. TABLE S4**.

**TABLE 2:** The top 5 most abundant genera for each of the three *Xenopus* sample types (based on mean relative abundance).

| Sample Type | Genus | Phylum | Abundance |
| --- | --- | --- | --- |
| Mother skin swab | *Chryseobacterium* | Bacteroidota | 0.27 |
|  | *Vogesella* | Proteobacteria | 0.19 |
|  | *Acinetobacter* | Proteobacteria | 0.12 |
|  | *Pseudomonas* | Proteobacteria | 0.05 |
|  | *Rheinheimera* | Proteobacteria | 0.04 |
| Egg jelly wash | *Chryseobacterium* | Bacteroidota | 0.12 |
|  | *Acinetobacter* | Proteobacteria | 0.12 |
|  | *Aeromonas* | Proteobacteria | 0.08 |
|  | *Lachnospiraceae NK4A136* | Firmicutes | 0.06 |
|  | *Bacteroides* | Bacteroidota | 0.04 |
| Tadpole tail | *Rhizobium* Genera | Proteobacteria | 0.38 |
|  | *Chryseobacterium* | Bacteroidota | 0.24 |
|  | *Fluviicola* | Bacteroidota | 0.07 |
|  | *Aeromonas* | Proteobacteria | 0.04 |
|  | *Delftia* | Proteobacteria | 0.04 |

### 3.2 The influence of the environment on *Xenopus* microbiomes is dependent on life stage

To assess the variability of microbial communities within the various sample types, Bray-Curtis distances were compared between samples of the same type (“within-type variation”) and between samples of different types (“between-type variation”). A generally high degree of within-type variation was observed for all sample types, with frog samples (mother skin swabs, egg jelly wash and tadpole tail clippings) generally being more variable than environmental samples (**FIGURE 3A, SUPP. TABLE S5A-C**).

**Figure 3.**
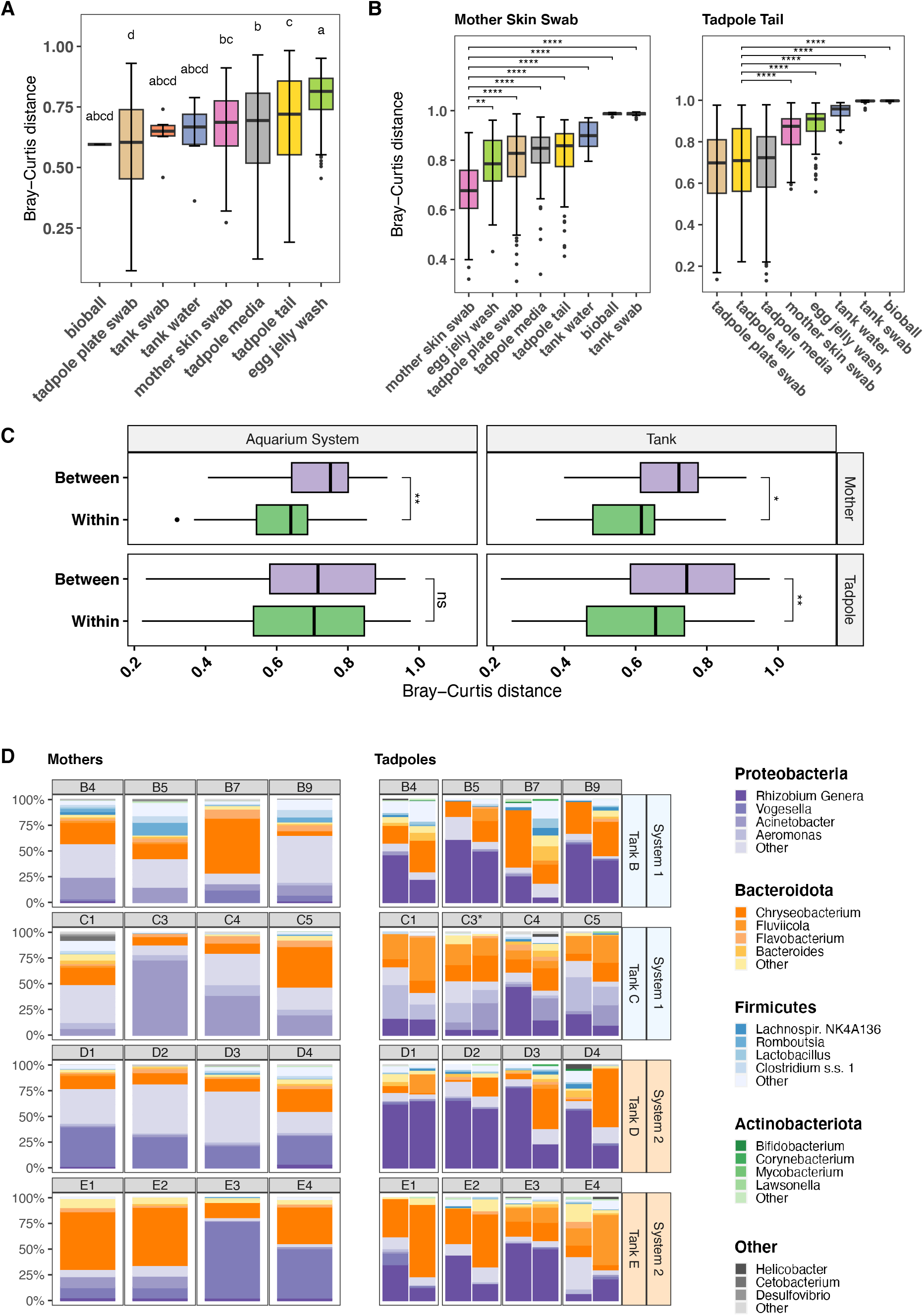
Variability of microbial communities in core sample types based on 16S amplicon sequencing. A) Box plot of Bray-Curtis distances between samples of the same type. Each box shows the distribution of distances between samples of that sample type. Bars within boxes indicate median value. Statistical significance is shown using compact letter display, sample types within the same letter group have no statistically significant difference from each other based on Dunn’s post-hoc test (p < 0.05). B) Box plot showing relative distances between mother skin swab samples and tadpole tail samples and other sample types. Asterisks indicate significance levels determined by Dunn’s post-hoc test; **** = < 0.0001, *** = < 0.001, ** = < 0.01, * = < 0.05. Values are not shown for pairs with non-significant p-values. C) Comparison of Bray-Curtis distance variation in mother skin swabs and tadpole tail samples. On the Y axis, ‘Within’ refers to distances between samples from the same tank or from within the same aquarium system according to the panel. ‘Between’ refers to distances between samples taken from different tanks or systems. Asterisks indicate significance levels determined by Wilcoxon Rank Sum; **** = < 0.0001, *** = < 0.001, ** = < 0.01, * = < 0.05, ns = not significant. D) Comparison of relative abundance of microbial phyla and genera on individual mother skin and tadpole tail samples within sibships. Each bar within the tadpole panel represents tadpoles from an individual plate (i.e. two plates per mother). Samples are rarefied at 1000 reads, with the exception of the first plate of C3 tadpoles (denoted with a *); for this plate there were insufficient reads to rarefy at 1000, so abundance is based on the unrarefied reads (650 mean reads). Horizontal labels indicate identity of mother, while vertical labels on right of plot indicate mother’s tank of residence and the aquarium system the tank is within (A or B). Mother samples represent multiple skin swabs, while tadpole samples are tail clips from multiple tadpoles per mother.

Between-type distances were frequently greater than within-type distances. We focussed on the distances between mother skin swab or tadpole tail samples and other sample types (**FIGURE 3B**). For mother skin swabs, the greatest median distances were observed to tank samples, indicating little relationship between the mother skin microbiome and their environment; distances between mother samples were significantly shorter than between mothers and other sample types (**SUPP. TABLE S6A-C**). For tadpole tail samples, distances to corresponding environmental samples (tadpole media/plate swabs) were similar to tadpole tail within-type distances (**SUPP. TABLE S6A-C**), suggesting that tail microbiomes are more similar to their environment.

We also calculated and compared mean pairwise Pearson’s correlation coefficients within and between all sample types (**SUPP. FIGURE S4**). Samples were strongly correlated with other samples of the same type and less so with samples of different types, although tadpole tail samples were also strongly correlated with tadpole environmental samples.

To determine whether the mother’s tank or aquarium system influenced the mother or tadpole skin microbiome, Bray-Curtis distances between samples within the same tank or aquarium system were compared with distances between samples from different tanks or aquaria (**FIGURE 3C**) (refer to **FIGURE 1, SUPP. FIGURE S1** and **SECTION 2.2** for detail on tank/aquarium system structure). For mother skin microbiome samples, there was significantly less distance between samples from the same tank or aquarium than between those from different tanks/aquaria (Wilcoxon Rank Sum, **SUPP. TABLE S7B)**, indicating that both variables affect the adult skin microbiome. PERMANOVAs confirmed a significant effect of aquarium system, which accounted for 25% of variation observed, while the interaction between aquarium system and tank was not significant (**SUPP. TABLE S7C**). For tadpoles, there was a significant difference observed between within- and between-group distances for tanks, but not for aquarium system (Wilcoxon Rank Sum, **SUPP. TABLE S7B**). The aquarium system accounted for 8% of variation, whereas the combined effect of aquarium system and tank accounted for 20.8%. (PERMANOVA, **SUPP. TABLE S7C**). A second PERMANOVA to assess the effect of tank and female found that the tank accounted for 28% of variation, while the combined effect of tank and mother ID accounted for 41% (**SUPP. TABLE S7C**).

The composition of microbiomes from each female and each plate of tadpoles within a sibship are shown in **FIGURE 3D**. Differences in mother skin microbiome can be seen according to tank as well as aquarium system. *Vogesella* was more abundant in mothers from System 2 (Tanks D and E), while *Acinetobacter* was proportionately more common in System 1 (Tanks B and C). *Chryseobacterium* (followed by *Vogesella*) was the overall most abundant genus in Tank E, although *Vogesella* was more common on two frogs. Firmicutes did not comprise a majority of microbes from any frog, but they were more common in frogs from Tank B (mean 18%, c.f. 1 – 7%), which comprised frogs considerably older than those from Tanks C – E (**SUPP. TABLE S8**). Tadpole tail microbiome composition also appeared to vary according to the mother’s tank; for example, tadpoles from Tank D mothers were most often dominated by *Rhizobium*, which was in relatively low abundance in Tank C sibships. Firmicutes were in highest abundance in Tank B tadpoles, which corresponds to a higher profusion of this phylum in Tank B mother frogs.

### 3.3 Tadpole skin microbiomes are similar to their environment, while those of adult frogs are more distinct

Differential abundance of taxa between mother skin swab and tadpole tail samples was assessed using ANCOMBC2 and MaAsLin2. Thirty-four taxa were found to be significantly differentially abundant (alpha = 0.25) by both analyses, with a further 21 taxa identified by one analysis but not the other. The mean relative abundances of the 34 common taxa are shown for each of the sample types in **FIGURE 4A**; full results of the analyses can be found in **SUPP. TABLES S9 & S10**. Further, the mean relative abundance across sample types of a selection of taxa that were notably abundant in **SUPP. TABLE S4** are shown in **FIGURE 4B**.

**Figure 4.**
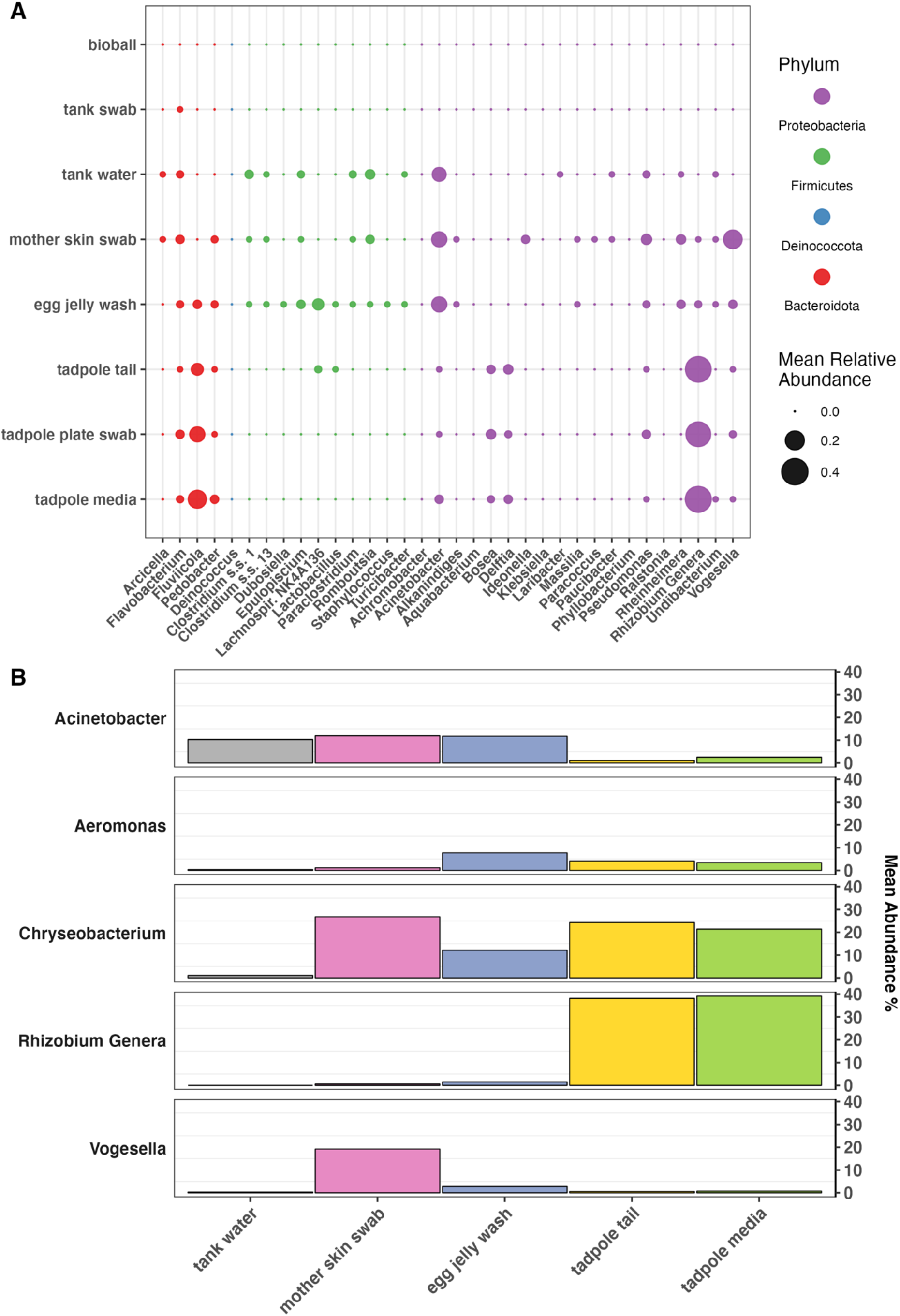
A) Mean relative abundance of significantly differentially abundant taxa across sample types. Taxa shown here are those found to be differentially abundant between mother skin swabs and tadpole tails by both ANCOMBC2 and MaAsLin2 (q < 0.25). B) Mean relative abundance of selected taxa across sample types, demonstrating changes in abundance through the progression of the *X. laevis* life cycle (left to right). Taxa shown were noted as being particularly abundant across one or more sample types of interest in **SUPP. TABLE S4**.

*Rhizobium* were markedly more abundant in tadpole tails and their immediate environment than any other sample type, while taxa such as *Romboutsia, Acinetobacter* and *Clostridium* were more abundant in the tank environment and/or on mothers and eggs (**FIGURES 4A & 4B**). Meanwhile, *Vogesella* was notably abundant on mother frogs and to a lesser extent, eggs, while being relatively uncommon in the corresponding environment. Mother and tank water samples seemed to be less favourable to taxa such as *Fluviicola* and *Rhizobium*. *Chryseobacterium* was abundant in all three frog life stages, as well as in tadpole media.

A microbial source tracking analysis was completed using FEAST v0.1.0 (**FIGURE 5; SUPP. TABLE S11**). While the source of a proportion of microbial taxa was ultimately unknown for each sink (mother skin, egg jelly, and tadpole environmental samples) the analysis indicates the likely most significant sources of microbiomes. The skin microbiome of female adult frogs was largely sourced from their tank water, while the bacteria adhering to the tank wall made an almost negligible contribution. The majority of the microbiome associated with egg jelly was sourced from the mother’s skin, and tadpoles subsequently received most of their skin microbiome from the eggs before seeding their environment (MMR media).

**Figure 5.**
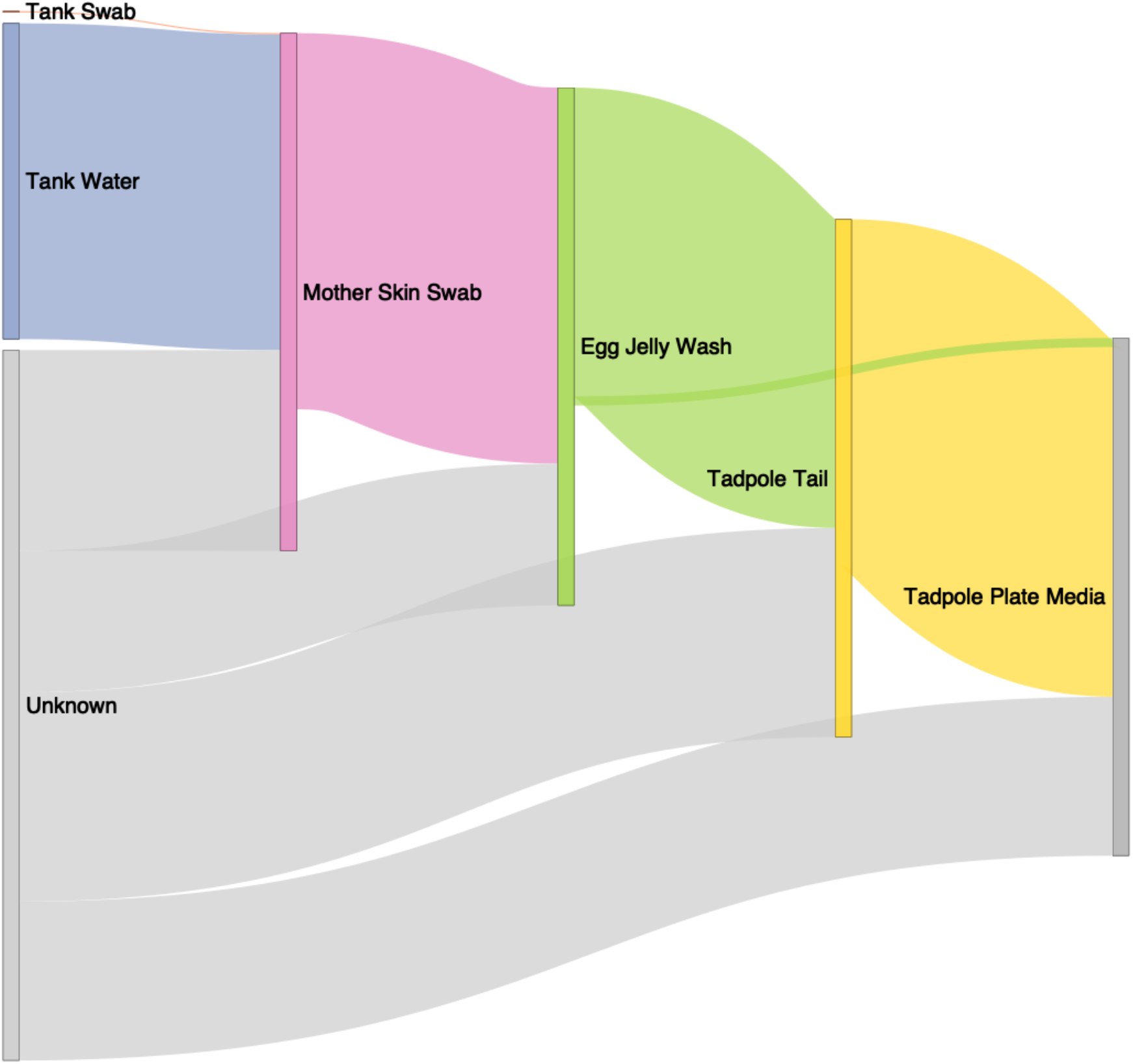
Sankey diagram conceptualising flow of microbial taxa through the environment and frog life stages, based on FEAST analysis. Bioballs are not included in this analysis as they are not in direct contact with frogs and are similar in composition to tank wall swabs. Tadpole plate swabs were not analysed as a sink due to their similar composition to tadpole plate media.

## 4 Discussion

Frogs of the *Xenopus* genus, most notably *X. laevis* and *X. tropicalis*, are widely used model organisms, finding particular utility in tissue regeneration, developmental, genetic, and neurobiological research. Many frogs are supplied to research laboratories by central breeding facilities which routinely treat tadpoles with antibiotics (e.g. gentamicin) and/or thimerosal [60, 61] which inherently alters microbiome community structure and transmission [30]. However, frogs are sometimes also raised in smaller, in-house facilities which may employ different rearing protocols, which, in combination with local environmental differences, is likely to affect microbiome composition. It is becoming evident that bacteria play an important role in *Xenopus* tail regeneration [29, 30] and are undoubtedly critical in the development of immune and other physiological systems. For this reason, it is imperative to characterise microbiomes associated with *Xenopus* and consider how they may be influencing experimental results.

A large proportion of studies into amphibian skin microbiota to date have been undertaken with reference to chytridiomycosis (summarised by Jiménez & Sommer [62]). It is demonstrated that chytrid infection drives changes in the skin microbiome [14, 63–66]; indeed, some bacterial taxa are shown to have anti-chytrid properties [67, 68]. These taxa are conceivably selected for in epi/enzootic scenarios, and have been investigated as potential probiotic therapies [68–70]. The *X. laevis* colony involved in this study has previously tested negative to chytrid infection, and is assumed to remain so due to the closed nature of the colony, biosecurity protocols and lack of symptoms. However, we note the presence of potentially anti-fungal genera identified by Woodhams et al. [68], such as *Chryseobacterium* and *Pseudomonas*, on our frogs and tadpoles.

### Mother *Xenopus* skin microbiomes are affected by the environment, but the frogs have ultimate control

The most complex microbiomes in our study were observed in tank environmental samples (water, wall swabs and bioballs) while the least complex were tank media and plate swabs. The tank environment is well established and has potential inputs from a range of sources including tap water, water conditioning additives, frog food, human attendants, and the frogs themselves, while the microbiomes of tadpole environment samples are at an early stage of establishment and limited to those introduced by the tadpole itself or incidental/airborne contaminants. Mother frogs’ skin microbial composition was the most complex of the frog derived samples, however was less diverse than their environment; indeed, mother frogs had the highest ‘unknown’ microbiome source of any life stage, which suggest they may be selecting for microbes occurring at low abundance in the environment (e.g. *Vogesella*). Tank water samples were closer in distance to mothers than were tank wall or bioball samples. We note that although mothers were moved into clean filtered water before swabbing, they were not rinsed to remove transient environmental DNA, which may have resulted in some microbiota remaining on the skin. However, tank and mother sample analyses suggest that the adult *X. laevis* skin favours certain skin microbiota adapted to withstand and exploit the unique physical conditions. This is a similar conclusion to Piccinni et al. [34]’s laboratory *X. laevis* study, and was also suggested by Li et al. [33]’s research on *X. tropicalis* froglet gut microbiomes. Studies of wild amphibians [18, 19, 71, 72] yield comparable results. Host control of the skin microbiome could be accomplished through a variety of mechanisms, including skin shedding [73–75] and mucous secretion [19]. Importantly, amphibian skin mucous unique assemblages of anti-microbial peptides [19, 68, 76–78] that probably dictate the suite of microbes able to successfully colonise. The microbes themselves produce metabolites that may inhibit growth of other microbes [11–13]. While female frogs are likely to select for certain skin microbiota through the above means, it was clear from the analysis of Bray-Curtis distances that microbiomes still vary between individual frogs, and are also affected by the tank and aquarium system. Individual differences in physiology and immune development may account for some variation between frogs. Frogs from Tank B were born in 2007, while those from other tanks were considerably younger (2016 – 2018), so age-related physiological factors may account for some variation observed in this study. Differences between tanks may also be attributed to small differences in environmental factors due to the tank’s location in the aquarium room, or may be related to the cohort of tank mates and the assemblage of microbiota each potentially shares with others when introduced. The two aquarium systems, located on opposite sides of the room, may also be subject to microenvironmental differences as for tanks, as well as subtle differences in filtration, water flow, etc. which could impact microbiota.

### Mother skin seeds the egg jelly, which in turn provides some microbiota to the tadpoles

A significant portion of the tadpole skin microbiome appears to be received vertically from the mother via the egg jelly. A large portion (mean 69.8% +/- 20.4) of the egg jelly microbiome was derived from the mother’s skin. Scalvenzi et al. [32] reported that, while in many cases a large portion of the *X. tropicalis* egg microbiome could not be traced to the parents, the parent’s skin contributed significantly more than did their faeces/gut. The relative unimportance of the adult gut microbiome in seeding egg microbiomes is surprising given the close contact between eggs and the female digestive tract in the amphibian cloaca. We did not sample the parent gut microbiome in this study as this was difficult to achieve effectively without sacrificing female frogs, however the skin appears to be a major contributor to egg microbial flora. Scalvenzi et al. [32]’s study differs from the present experiment in that parents were allowed to mate naturally, whereas our embryos were produced using in vitro fertilization. Eggs therefore had no direct contact with male frogs. While testis samples were sequenced and returned variable read counts, a large proportion had low bacterial biomass, and it was deemed likely that contamination during the necropsy and collection process was responsible for most reads. Testes are therefore considered unlikely to be a primary contributor to colonisation of eggs and subsequent life stages under the study conditions. Given that the majority of laboratories produce *Xenopus* embryos using in vitro fertilization, the mother is logically a more significant source of microbes in these cases.

Kueneman et al. found that larval stages of boreal toads (*Anaxyrus boreas*) [72] and Cascades frog (*Rana cascadae*) [71] had the lowest skin microbiome alpha diversity of all life stages, and we demonstrated the same for *X. laevis*. We also found that tadpole microbiomes were more variable than their mothers; this may be at least partially due to tadpole replicates representing different individuals rather than true technical replicates. However, newly developed embryos and tadpoles are also a prospect for colonisation by opportunistic microbiota, and their early communities may reflect those taxa which either happen to encounter them first or are most suited to rapid exploitation of that environment. McGrath- Blaser et al. [79] determined that tadpole microbiomes established via both vertical transmission and environmental exposure in Bornean foam-nesting frogs (*Polypedates* spp.). The sibship (mother ID), mother’s tank and aquarium system were observed to affect tadpole microbiomes, and these effects may be directly passed vertically to tadpoles from the mothers via eggs. While a proportion of our tadpole microbiota were contributed by the egg jelly, this leaves a proportion derived from unknown sources. It was not considered likely that the plates (petri dishes) or media (0.1x MMR) were significant contributors. Plates used were new and provided in sterile packaging, and media was autoclaved prior to use. While this should have minimized the number of microbes present, it is possible that some bacteria may have entered media bottles after opening. Media samples sequenced here (as well as plate swabs) were collected after exposure to tadpoles for a period of one week, and are quite similar in microbial composition to tadpole samples. We determined that, given the high calculated contribution of the egg jelly to tadpole samples and previous records of dominant taxa in frogs/tadpoles, it was likely that tadpoles were seeding their environment rather than vice versa. However, in future it would be preferable to sequence the media prior to tadpole exposure.

At each stage of the *X. laevis* life cycle, a notable proportion of the microbiome was attributed to unknown sources. We used a variety of controls to detect potential contaminants introduced throughout the sample collection and processing pipeline. The ‘kit microbiome’ or ‘kitome’ is a recognised source of contaminants during DNA extraction [80] and while we cannot eliminate this as a contributor to ‘unknown’ sources, our blank controls revealed negligible reads and we have taken measures to computationally account for and remove contaminants prior to analysis. Other potential ‘unknown’ sources include airborne microbiota in the lab and those potentially inadvertently introduced by the sample collector.

### Proteobacteria and Bacteroidota dominate the *X. laevis* skin microbiome

Proteobacteria and Bacteroidota were the dominant phyla across frog, egg and tadpole samples. This is similar to Piccinni et al. [34]’s study which found that Bacteroidota (in particular Flavobacteriia) followed by Proteobacteria were the most common bacterial phylum on adult *X. laevis*. Proteobacteria were dominant in their feeding stage tadpoles, with Bacteroidota making up a relatively minor portion of the skin microbiome.

In our study, tadpole skin microbiomes were dominated by a genus of Proteobacterium, *Rhizobium* (designated as *Allorhizobium-Neorhizobium-Pararhizobium-Rhizobium* by the SILVA v. 138.1 database). Most *Rhizobium* are gram-negative, nitrogen fixing plant symbionts and would not be immediately associated with the aquatic environment; however some species are free-living [81]. While *Rhizobium* were not among the top 50 taxa in Piccinni et al. [34]’s *X. laevis* microbiome dataset, bacteria from the Rhizobiales order were reported from Scalvenzi et al. [32]’s *X. tropicalis* metamorph/froglet guts, and Rhizobiaceae were detected in post-metamorphic *X. laevis* by Mashoof et al. [31]. A *Rhizobium* genus was also associated with faeces and gut samples in laboratory raised Asiatic toad tadpoles (*Bufo gargarizans*) [82]. Interestingly, a previous study of tadpoles from the same colony as this study Chapman et al. [30] did not detect this *Rhizobium* genotype, nor did it identify any *Rhizobium* as a dominant genus despite an absence of major changes in husbandry. These tadpoles were instead dominated by *Shinella* or *Delftia* (for antibiotic treated tadpoles) or *Shinella, Acinetobacter, Escherichia-Shigella* or *Chryseobacterium* (untreated). The differences between the two studies may be attributed to different sampling techniques (Chapman et al. [30] extracted DNA from the entire tadpole, while the current study used only tail clippings), antibiotic effects in the earlier study, fluctuations in microbe populations in the various inputs (food, water etc.) or natural community drift/succession. Differences in computational processing and analysis of data may also be a factor.

The other notably dominant taxa on both mothers and tadpoles was a genus of Bacteroidota, *Chryseobacterium* (once classified as *Flaviobacterium* [83]). In some tadpoles, this taxa was more abundant than *Rhizobium*, and was also relatively common in tadpoles from the Chapman et al. [30] study. Neither Scalvenzi et al. [32] nor Piccinni et al. [34] reported *Chryseobacterium* from their tadpoles, however several epibiotic strains of ultra-small *Chryseobacterium* have previously been reported from adult *X. laevis* [84]. Piccinni et al. [34] did report that *Bergeyella*, a related species [83, 85] was dominant on adult *X. laevis* skin, although very low or undetectable in tadpole and water samples. *Chryseobacterium* is also well documented from newts [86] and fish [87] and the closely related *Elizabethkingi* genus (once considered to be a *Chryseobacterium*) has been associated with disease in frogs [88–90] including *X. laevis* [91]. We have successfully cultured *Chryseobacterium* from *X. laevis* adults and tadpoles and found that it correlated with improved tadpole tail regeneration outcomes [30].

Other notable taxa on frog samples included *Acinetobacter*, *Aeromonas*, *Fluviicola*, and *Vogesella*. A strain of the latter genus (*Vogesella* sp. Strain XCS3) was successfully cultured and sequenced from our frogs by Hudson et al. [92]. Tank/aquarium samples were dominated by both Proteobacteria and Bacteroidota as well as Acidobacteriota. *Aeromonas*, a taxon commonly found in aquatic environments and among the most abundant in our egg jelly samples, was only reported by Piccinni et al. [34] on ‘clean’ (thimerosal/ethanol treated) tadpoles and in standard ‘unclean’ water, while the remaining taxa listed above were not within their top 50 abundant taxa. *Aeromonas* was common in Scalvenzi et al. [32]’s tadpole gut samples, but the others were once again absent.

### Captive *X. laevis* microbiomes depend on laboratory environment and husbandry

As is clear from the above, microbiomes appear to vary between species of the same host genus, colonies, tanks/aquaria and probably over time. The marked differences are likely due to differences in skin microenvironment (skin structure, secretions, etc.) between species, variations in environment (i.e. microbial inputs from water and surrounding environment) and husbandry (e.g. food source, aquarium additives, antibiotic use). We note that Piccinni et al. [34]’s tadpoles were approximately one month old, pre-metamorphic, and being fed on algae. We sampled our tadpoles at a stage where they were not sufficiently developed to require formal feeding (stage 46, [35]), as food input may have confounded attempts to detect vertical transmission of microbes. Any potential food intake for our tadpoles would have been comprised of material derived from other tadpoles (tissue or waste product) or other incidental microscopic items scavenged from the media, with waste material derived from these sources or from yolk digestion/absorption. The difference in feeding status is likely to impact opportunities for colonisation of the gut and the skin through different/additional microbial inputs into the tadpole environment, and through concurrent changes to the gut and immune system associated with development stage. In our colony, we determined that frog food (generic salmon pellets) was unlikely to contribute substantively to the skin microbiome of adult frogs due to a low number of bacterial reads; we also determined that the tank water conditioner, comprised almost entirely of Firmicutes, was not a major contributor.

## 5 Conclusions

This study has explored the microbial communities associated with non-antibiotic treated, captive *X. laevis* and the dynamics of transmission between different frog life stages and their environment. Mother frog skin microbiomes were less diverse than those in their environment, and were dominated by different taxa, indicating that adult frogs have some ability to select for certain taxa that are present at low abundances in their environment. Tadpoles inherit a proportion of their microbiomes from their mothers via the egg, however have a less diverse microbial community that is distinct from their parents, which may reflect opportunistic colonisation of an available niche. After comparison with previous studies, these results indicate that although adult frogs retain some control over their microbiomes, the ultimate composition is dependent on environmental and experimental conditions. The microbiome and the impact of husbandry procedures should be an important consideration in *Xenopus* laboratory experiments. In lieu of microbial community profiling, detailed reporting of husbandry practices and environmental conditions should accompany all experimental results.

## 6 Declarations

### 6.1 Ethics approval and consent to participate

*X. laevis* egg and embryo production methodology was approved by the University of Otago’s Animal Ethics Committee (permit no. AUP19-01).

### 6.2 Consent for publication

Not applicable

### 6.3 Availability of data and material

R code for all analyses is provided at https://gitlab.com/pachapman/xenopus_microbiomes. Sequence data has been deposited in Sequence Read Archive (SRA) under BioProject ID PRJNA823390 (BioSamples SAMN33958523 - SAMN33958729).

### 6.6 Author’s contributions

XCM and CWB conceived and designed the study. DH and CWB conducted the experiments. PAC and XCM performed the analyses. PAC, XCM and CWB contributed to the preparation and editing of the manuscript and figures. All authors read and approved the final manuscript.

## Acknowledgements

We would like to acknowledge Joanna Ward and Nikita Woodhead for their skilled assistance in the laboratory and the aquarium respectively.

