## Supplementary_data for "The role of family and environment in determining the skin microbiome of captive aquatic frogs, *Xenopus laevis*": Xenoflow_supps_R4.html

Supplementary Data: Chapman et al. 2023


#### Contents

- Figure S1: Sampling Structure
- Table S1: Read Summary Statistics
- Figure S2A & S2B: Alpha Diversity
  - A. Shapiro Wilk tests of normality
  - B. Kruskal-Wallis tests
  - C. Dunn’s post-hoc tests
- **Table S3: Fig. 2B PERMANOVA**
- **Figure S3: Fig. 2B Silhouette analysis**
- Table S4: Fig. 2C Top 5 taxa
- Table S5A-C: Figure 3A supporting statistics
  - A. Shapiro Wilk test of normality
  - B. Kruskal-Wallis test
  - C. Dunn’s post-hoc test
- Table S6A-C: Figure 3B supporting statistics
  - A. Shapiro Wilk tests of normality
  - B. Kruskal-Wallis tests
  - C. Dunn’s post-hoc tests
- Figure S4: Pearson’s correlation coefficients between sample types
- Table S7A-C: Figure 3C supporting statistics
  - A. Shapiro Wilk test of normality
  - B. Wilcoxon Rank Sum tests
  - C. PERMANOVAs
- **Table S8: Abundance of phyla on mother skin swab samples**
- **Table S9: ANCOMBC2 results**
- **Table S10: MaAsLin2 results**
- Table S11: FEAST results summary

### Supplementary Data: Chapman et al. 2023

Authors

Phoebe A. Chapman

Daniel Hudson

Xochitl C. Morgan

Caroline W. Beck

### Figure S1: Sampling Structure

****FIGURE** **S1:****  sampling strategy. Numbers within icons on left panel denote numbers of biological replicates collected per sample type, with technical replicates indicated within accompanying text.

### Table S1: Read Summary Statistics

| **Sample Type** | Samples | Min | Median | Mean | Max | SD |
| --- | --- | --- | --- | --- | --- | --- |
| bioball | 2 | 2141.50 | 2203.00 | 2203.00 | 2264.50 | 86.97 |
| control Tween | 1 | 0.50 | 0.50 | 0.50 | 0.50 | NA |
| control extraction | 1 | 1.00 | 1.00 | 1.00 | 1.00 | NA |
| control filter | 1 | 0.50 | 0.50 | 0.50 | 0.50 | NA |
| control mock | 1 | 2811.00 | 2811.00 | 2811.00 | 2811.00 | NA |
| control swab | 1 | 0.00 | 0.00 | 0.00 | 0.00 | NA |
| egg jelly wash | 16 | 793.00 | 4142.75 | 3991.94 | 7879.50 | 2113.89 |
| frog food | 1 | 584.00 | 584.00 | 584.00 | 584.00 | NA |
| mother skin swab | 16 | 155.50 | 2985.25 | 4222.62 | 10270.00 | 3212.31 |
| stresszyme | 1 | 5612.00 | 5612.00 | 5612.00 | 5612.00 | NA |
| tadpole media | 32 | 1087.00 | 3947.50 | 4384.16 | 13201.00 | 2949.63 |
| tadpole plate swab | 32 | 1087.50 | 5599.50 | 6642.78 | 13870.50 | 3364.99 |
| tadpole tail | 32 | 586.50 | 3853.25 | 5532.92 | 15664.50 | 4548.66 |
| tank swab | 4 | 1261.50 | 2528.00 | 2657.62 | 4313.00 | 1581.47 |
| tank water | 4 | 1236.00 | 3318.00 | 3592.62 | 6498.50 | 2176.33 |
| testis | 4 | 289.00 | 2473.25 | 3134.88 | 7304.00 | 3215.49 |
|  |  |  |  |  |  |  |
| --- | --- | --- | --- | --- | --- | --- |
| **TABLE S1:** Read count summary statistics, reported per sample type. Samples - total number of samples of that type following merging of replicates; Min - minimum; Max - maximum; SD - standard deviation | | | | | | |

### Figure S2A & S2B: Alpha Diversity

****FIGURE S2:****  plots summarising alpha diversity metrics for core sample types. A. Shannon Diversity B. Faith’s Phylogentic Diversity. Statistical significance is shown using compact letter display, sample types within the same letter group have no statistically significant difference from each other based on Dunn’s post-hoc test (p < 0.05)

#### A. Shapiro Wilk tests of normality

|  | W statistic | p-value |
| --- | --- | --- |
| Observed | 0.80 | 5.45 × 10^−12^ |
| Shannon | 0.95 | 4.57 × 10^−5^ |
| Faith's Phylogenetic | 0.78 | 9.57 × 10^−13^ |
|  |  |  |
| --- | --- | --- |
| **TABLE S2A:** Results of Shapiro Wilk tests of normality on three alpha diversity metrics. Red text indicates a statistically significant p-value. A signficant p-value indicates a non-normal distribution. p < 0.05 | | |

#### B. Kruskal-Wallis tests

|  | KW chi-squared (H) | df | p-value |
| --- | --- | --- | --- |
| Observed | 89.54 | 7 | 2.20 × 10^−16^ |
| Shannon | 67.37 | 7 | 5.02 × 10^−12^ |
| Faith's Phylogenetic | 92.29 | 7 | 2.20 × 10^−16^ |
|  |  |  |  |
| --- | --- | --- | --- |
| **TABLE S2B:** Results of Kruskal-Wallis tests comparing sample types for three alpha diversity metrics. Red text indicates a statistically significant p-value. df = degrees of freedom. p < 0.05 | | | |

#### C. Dunn’s post-hoc tests

| Comparison | **Observed** | | **Shannon** | | **Faith’s Phylogenetic** | |
| --- | --- | --- | --- | --- | --- | --- |
|  | Z | p.adj | Z | p.adj | Z | p.adj |
| bioball - egg jelly wash | 0.91 | 1.00 | 0.65 | 1.00 | 0.90 | 1.00 |
| bioball - mother skin swab | 0.94 | 1.00 | 1.40 | 1.00 | 1.08 | 1.00 |
| bioball - tadpole media | 3.21 | 3.66 × 10^−2^ | 3.02 | 7.00 × 10^−2^ | 3.05 | 6.34 × 10^−2^ |
| bioball - tadpole plate swab | 3.35 | 2.24 × 10^−2^ | 2.88 | 1.11 × 10^−1^ | 3.55 | 1.07 × 10^−2^ |
| bioball - tadpole tail | 1.87 | 1.00 | 1.99 | 1.00 | 1.81 | 1.00 |
| bioball - tank swab | 0.29 | 1.00 | 0.11 | 1.00 | 0.16 | 1.00 |
| bioball - tank water | −0.06 | 1.00 | −0.04 | 1.00 | −0.05 | 1.00 |
| egg jelly wash - mother skin swab | 0.06 | 1.00 | 1.55 | 1.00 | 0.38 | 1.00 |
| egg jelly wash - tadpole media | 5.29 | 3.39 × 10^−6^ | 5.48 | 1.18 × 10^−6^ | 4.95 | 2.04 × 10^−5^ |
| egg jelly wash - tadpole plate swab | 5.61 | 5.55 × 10^−7^ | 5.15 | 7.37 × 10^−6^ | 6.11 | 2.74 × 10^−8^ |
| egg jelly wash - tadpole tail | 2.14 | 9.03 × 10^−1^ | 3.03 | 6.78 × 10^−2^ | 2.04 | 1.00 |
| egg jelly wash - tank swab | −0.77 | 1.00 | −0.69 | 1.00 | −0.96 | 1.00 |
| egg jelly wash - tank water | −1.31 | 1.00 | −0.93 | 1.00 | −1.27 | 1.00 |
| mother skin swab - tadpole media | 5.22 | 4.95 × 10^−6^ | 3.68 | 6.60 × 10^−3^ | 4.51 | 1.83 × 10^−4^ |
| mother skin swab - tadpole plate swab | 5.54 | 8.29 × 10^−7^ | 3.34 | 2.32 × 10^−2^ | 5.67 | 4.06 × 10^−7^ |
| mother skin swab - tadpole tail | 2.07 | 1.00 | 1.27 | 1.00 | 1.60 | 1.00 |
| mother skin swab - tank swab | −0.81 | 1.00 | −1.70 | 1.00 | −1.20 | 1.00 |
| mother skin swab - tank water | −1.35 | 1.00 | −1.93 | 1.00 | −1.52 | 1.00 |
| tadpole media - tadpole plate swab | 0.40 | 1.00 | −0.42 | 1.00 | 1.45 | 1.00 |
| tadpole media - tadpole tail | −3.75 | 4.93 × 10^−3^ | −2.88 | 1.12 × 10^−1^ | −3.47 | 1.46 × 10^−2^ |
| tadpole media - tank swab | −3.94 | 2.25 × 10^−3^ | −3.97 | 2.01 × 10^−3^ | −3.94 | 2.30 × 10^−3^ |
| tadpole media - tank water | −4.51 | 1.81 × 10^−4^ | −4.22 | 6.95 × 10^−4^ | −4.27 | 5.46 × 10^−4^ |
| tadpole plate swab - tadpole tail | −4.14 | 9.72 × 10^−4^ | −2.47 | 3.73 × 10^−1^ | −4.87 | 3.11 × 10^−5^ |
| tadpole plate swab - tank swab | −4.13 | 1.00 × 10^−3^ | −3.77 | 4.52 × 10^−3^ | −4.62 | 1.07 × 10^−4^ |
| tadpole plate swab - tank water | −4.70 | 7.27 × 10^−5^ | −4.02 | 1.63 × 10^−3^ | −4.95 | 2.03 × 10^−5^ |
| tadpole tail - tank swab | −2.10 | 1.00 | −2.55 | 3.06 × 10^−1^ | −2.23 | 7.26 × 10^−1^ |
| tadpole tail - tank water | −2.66 | 2.19 × 10^−1^ | −2.79 | 1.48 × 10^−1^ | −2.56 | 2.95 × 10^−1^ |
| tank swab - tank water | −0.43 | 1.00 | −0.18 | 1.00 | −0.25 | 1.00 |
|  |  |  |  |  |  |  |
| --- | --- | --- | --- | --- | --- | --- |
| **TABLE S2C:** Results of Dunn's post-hoc tests on three alpha diversity metrics. Red text indicates a statistically significant p-value. Z = Z test statistic. p.adj = p-value with Bonferroni adjustment. p < 0.05 | | | | | | |

### **Table S3: Fig. 2B PERMANOVA**

|  | Df | SumOfSqs | R^2^ | F | p-value |
| --- | --- | --- | --- | --- | --- |
| SampleType | 7 | 11.65 | 0.28 | 8.08 | 1.00 × 10^−3^ |
| System | 1 | 1.19 | 0.03 | 5.76 | 1.00 × 10^−3^ |
| System:Tank | 3 | 3.62 | 0.09 | 5.86 | 1.00 × 10^−3^ |
| Residual | 125 | 25.73 | 0.61 | NA | NA |
| Total | 136 | 42.19 | 1.00 | NA | NA |
|  |  |  |  |  |  |
| --- | --- | --- | --- | --- | --- |
| **TABLE S3:** Results of PERMANOVA (adonis2) comparing Bray-Curtis distances between samples of different types. Aquarium system and Tank were included as fixed effects with interaction. Red text indicates a statistically significant p-value. Df = degrees of freedom. SumofSqs = sum of squares. R2 = R2 statistic. F = F statistic. | | | | | |

### **Figure S3: Fig. 2B Silhouette analysis**

****FIGURE S3:**** Silhouette analysis to assess clustering of different sample types.

### Table S4: Fig. 2C Top 5 taxa

| **Genus** | **Phylum** | **Mean Relative Abundance** |
| --- | --- | --- |
| bioball | | |
| --- | --- | --- |
| JGI 0001001-H03 | Acidobacteriota | 0.24 |
| RB41 | Acidobacteriota | 0.11 |
| Piscinibacter | Proteobacteria | 0.06 |
| Rhizorhapis | Proteobacteria | 0.05 |
| Novosphingobium | Proteobacteria | 0.04 |
| egg jelly wash | | |
| Chryseobacterium | Bacteroidota | 0.12 |
| Acinetobacter | Proteobacteria | 0.12 |
| Aeromonas | Proteobacteria | 0.08 |
| Lachnospir. NK4A136 | Firmicutes | 0.06 |
| Bacteroides | Bacteroidota | 0.04 |
| mother skin swab | | |
| Chryseobacterium | Bacteroidota | 0.27 |
| Vogesella | Proteobacteria | 0.19 |
| Acinetobacter | Proteobacteria | 0.12 |
| Pseudomonas | Proteobacteria | 0.05 |
| Rheinheimera | Proteobacteria | 0.04 |
| tadpole media | | |
| Rhizobium Genera | Proteobacteria | 0.39 |
| Chryseobacterium | Bacteroidota | 0.21 |
| Fluviicola | Bacteroidota | 0.18 |
| Aeromonas | Proteobacteria | 0.03 |
| Pedobacter | Bacteroidota | 0.03 |
| tadpole plate swab | | |
| Rhizobium Genera | Proteobacteria | 0.35 |
| Chryseobacterium | Bacteroidota | 0.33 |
| Fluviicola | Bacteroidota | 0.12 |
| Bosea | Proteobacteria | 0.04 |
| Flavobacterium | Bacteroidota | 0.03 |
| tadpole tail | | |
| Rhizobium Genera | Proteobacteria | 0.38 |
| Chryseobacterium | Bacteroidota | 0.24 |
| Fluviicola | Bacteroidota | 0.07 |
| Aeromonas | Proteobacteria | 0.04 |
| Delftia | Proteobacteria | 0.04 |
| tank swab | | |
| Ferruginibacter | Bacteroidota | 0.11 |
| Hyphomicrobium | Proteobacteria | 0.10 |
| Stenotrophobacter | Acidobacteriota | 0.10 |
| Nitrospira | Nitrospirota | 0.08 |
| Rhizorhapis | Proteobacteria | 0.07 |
| tank water | | |
| Bacteroides | Bacteroidota | 0.12 |
| Cetobacterium | Fusobacteriota | 0.12 |
| Acinetobacter | Proteobacteria | 0.10 |
| Romboutsia | Firmicutes | 0.04 |
| dgA-11 gut group | Bacteroidota | 0.04 |
|  |  |  |
| --- | --- | --- |
| **Table S4**: The top 5 taxa from each of the core sample types, based on mean relative abundance | | |

### Table S5A-C: Figure 3A supporting statistics

#### A. Shapiro Wilk test of normality

| W statistic | p-value |
| --- | --- |
| 0.97 | 2.20 × 10^−16^ |
|  |  |
| --- | --- |
| **TABLE S5A:** Results of Shapiro Wilk tests of normality on Bray-Curtis distances. Red text indicates a statistically significant p-value. A signficant p-value indicates a non-normal distribution. p < 0.05 | |

#### B. Kruskal-Wallis test

| KW chi-squared (H) | df | p-value |
| --- | --- | --- |
| 157.75 | 7 | 2.20 × 10^−16^ |
|  |  |  |
| --- | --- | --- |
| **TABLE S5B:** Results of Kruskal-Wallis tests comparing Bray-Curtis distances within different sample types. Red text indicates a statistically significant p-value. df = degrees of freedom. p < 0.05 | | |

#### C. Dunn’s post-hoc test

| Comparison | Z | p.adj |
| --- | --- | --- |
| bioball - egg jelly wash | −1.24 | 1.00 |
| bioball - mother skin swab | −0.52 | 1.00 |
| bioball - tadpole media | −0.48 | 1.00 |
| bioball - tadpole plate swab | −0.12 | 1.00 |
| bioball - tadpole tail | −0.69 | 1.00 |
| bioball - tank swab | −0.19 | 1.00 |
| bioball - tank water | −0.27 | 1.00 |
| egg jelly wash - mother skin swab | 5.39 | 1.98 × 10^−6^ |
| egg jelly wash - tadpole media | 7.46 | 2.49 × 10^−12^ |
| egg jelly wash - tadpole plate swab | 11.06 | 5.79 × 10^−27^ |
| egg jelly wash - tadpole tail | 5.42 | 1.65 × 10^−6^ |
| egg jelly wash - tank swab | 2.47 | 3.82 × 10^−1^ |
| egg jelly wash - tank water | 2.28 | 6.37 × 10^−1^ |
| mother skin swab - tadpole media | 0.36 | 1.00 |
| mother skin swab - tadpole plate swab | 3.76 | 4.67 × 10^−3^ |
| mother skin swab - tadpole tail | −1.57 | 1.00 |
| mother skin swab - tank swab | 0.74 | 1.00 |
| mother skin swab - tank water | 0.55 | 1.00 |
| tadpole media - tadpole plate swab | 5.77 | 2.28 × 10^−7^ |
| tadpole media - tadpole tail | −3.26 | 3.12 × 10^−2^ |
| tadpole media - tank swab | 0.67 | 1.00 |
| tadpole media - tank water | 0.47 | 1.00 |
| tadpole plate swab - tadpole tail | −9.03 | 5.01 × 10^−18^ |
| tadpole plate swab - tank swab | −0.23 | 1.00 |
| tadpole plate swab - tank water | −0.42 | 1.00 |
| tadpole tail - tank swab | 1.17 | 1.00 |
| tadpole tail - tank water | 0.98 | 1.00 |
| tank swab - tank water | −0.14 | 1.00 |
|  |  |  |
| --- | --- | --- |
| **TABLE S5C:** Results of Dunn's post-hoc tests comparing Bray-Curtis distances within different sample types. Red text indicates a statistically significant p-value. z = Z statistic. p.adj = p-value with Bonferroni correction. p < 0.05 | | |

### Table S6A-C: Figure 3B supporting statistics

#### A. Shapiro Wilk tests of normality

|  | W statistic | p-value |
| --- | --- | --- |
| Mothers | 0.92 | 2.68 × 10^−15^ |
| Tadpoles | 0.92 | 2.68 × 10^−15^ |
|  |  |  |
| --- | --- | --- |
| **TABLE S5A:** Results of Shapiro Wilk tests of normality on Bray-Curtis distances. Red text indicates a statistically significant p-value. A signficant p-value indicates a non-normal distribution. p < 0.05 | | |

#### B. Kruskal-Wallis tests

|  | KW chi-squared (H) | df | p-value |
| --- | --- | --- | --- |
| Mothers | 106.33 | 7 | 2.20 × 10^−16^ |
| Tadpoles | 495.74 | 7 | 2.20 × 10^−16^ |
|  |  |  |  |
| --- | --- | --- | --- |
| **TABLE S6B:** Results of Kruskal-Wallis tests comparing Bray-Curtis distances within mother skin swab and tadpole tail samples and sample types. Red text indicates a statistically significant p-value. df = degrees of freedom. p < 0.05 | | | |

#### C. Dunn’s post-hoc tests

| Comparison | **Mothers** | | **Tadpoles** | |
| --- | --- | --- | --- | --- |
|  | Z | p.adj | Z | p.adj |
| bioball | 5.04 | 1.28 × 10^−5^ | 7.34 | 6.13 × 10^−12^ |
| egg jelly wash | 3.93 | 2.36 × 10^−3^ | 10.29 | 2.27 × 10^−23^ |
| mother skin swab | NA | NA | 7.58 | 9.69 × 10^−13^ |
| tadpole media | −6.42 | 3.81 × 10^−9^ | −0.26 | 1.00 |
| tadpole plate swab | −5.37 | 2.18 × 10^−6^ | −1.68 | 1.00 |
| tadpole tail | −6.61 | 1.05 × 10^−9^ | NA | NA |
| tank swab | −7.89 | 8.26 × 10^−14^ | −11.14 | 2.23 × 10^−27^ |
| tank water | −6.02 | 4.80 × 10^−8^ | −9.79 | 3.49 × 10^−21^ |
|  |  |  |  |  |
| --- | --- | --- | --- | --- |
| **TABLE S6C:** Results of Dunn's post-hoc tests comparing Bray-Curtis distances within and between mother skin swab and tadpole tail samples and other sample types. Red text indicates a statistically significant p-value. Z = Z test statistic. P (adj.) = p-value with Bonferroni adjustment. p < 0.05 | | | | |

### Figure S4: Pearson’s correlation coefficients between sample types

****FIGURE S4:**** Heatmap of mean Pearson’s correlation coefficients between samples of each core type.

### Table S7A-C: Figure 3C supporting statistics

#### A. Shapiro Wilk test of normality

|  | W statistic | p-value |
| --- | --- | --- |
| Mothers | 0.97 | 7.63 × 10^−3^ |
| Tadpoles | 0.95 | 5.95 × 10^−12^ |
|  |  |  |
| --- | --- | --- |
| **TABLE S7A:** Results of Shapiro Wilk tests of normality on Bray-Curtis distances in mother skin swab and tadpole tail samples. Red text indicates a statistically significant p-value. A signficant p-value indicates a non-normal distribution. p < 0.5 | | |

#### B. Wilcoxon Rank Sum tests

| Comparison | W statistic | p.adj |
| --- | --- | --- |
| mother skin swab | | |
| --- | --- | --- |
| Aquarium system | 531 | 4.32 × 10^−3^ |
| Tank | 372 | 2.59 × 10^−2^ |
| tadpole tail | | |
| Aquarium system | 8482 | 7.76 × 10^−1^ |
| Tank | 7115 | 2.33 × 10^−3^ |
|  |  |  |
| --- | --- | --- |
| **TABLE S7B:** Results of Wilcoxon Rank Sum tests comparing Bray-Curtis distances from mother skin swab and tadpole tail samples. Tests are comparing distances between samples from the same group (grouped by aquarium system or tank) with distances between samples from different groups. Red text indicates a statistically significant p-value. p.adj = p-value adjusted using Bonferroni correction; p < 0.05 | | |

#### C. PERMANOVAs

| **Comparison** | **Df** | **SumOfSqs** | R^**2**^ | **F** | **p-value** |
| --- | --- | --- | --- | --- | --- |
| mother skin swab | | | | | |
| --- | --- | --- | --- | --- | --- |
| System | 1.00 | 0.82 | 0.25 | 4.64 | 1.00 × 10^−3^ |
| System:Tank | 2.00 | 0.55 | 0.17 | 1.57 | 9.40 × 10^−2^ |
| Residual | 11.00 | 1.94 | 0.59 | NA | NA |
| Total | 14.00 | 3.31 | 1.00 | NA | NA |
| tadpole tail | | | | | |
| System | 1.00 | 0.61 | 0.08 | 2.97 | 2.10 × 10^−2^ |
| System:Tank | 2.00 | 1.67 | 0.21 | 4.07 | 2.00 × 10^−3^ |
| Residual | 28.00 | 5.73 | 0.72 | NA | NA |
| Total | 31.00 | 8.00 | 1.00 | NA | NA |
| Tank | 3.00 | 2.27 | 0.28 | 4.99 | 1.00 × 10^−3^ |
| Tank:Female | 12.00 | 3.30 | 0.41 | 1.81 | 9.00 × 10^−3^ |
| Residual | 16.00 | 2.43 | 0.30 | NA | NA |
| Total | 31.00 | 8.00 | 1.00 | NA | NA |
|  |  |  |  |  |  |
| --- | --- | --- | --- | --- | --- |
| **TABLE S7C:** Results of PERMANOVAs (adonis2) to test significance of differences in Bray-Curtis distances and the effect size of the mother's ID, tank, or aquarium system. For both sample types, aquarium system and tank were included as fixed effects with interaction. For tadpoles, a second PERMANOVA tested the effects of the mother's ID and tank separately. Red text indicates a statistically significant p-value. Df = degrees of freedom. SumofSqs = sum of squares. R2 = R2 statistic. F = F statistic. p < 0.05 | | | | | |

### **Table S8: Abundance of phyla on mother skin swab samples**

| **Tank** | **Mother Age** | **Actinobacteriota** | **Bacteroidota** | **Firmicutes** | **Proteobacteria** |
| --- | --- | --- | --- | --- | --- |
| B | 2007 | 0.01 | 0.32 | 0.18 | 0.48 |
| C | 2016 | 0.00 | 0.32 | 0.06 | 0.60 |
| D | 2018 | 0.00 | 0.22 | 0.07 | 0.69 |
| E | 2018 | 0.00 | 0.50 | 0.01 | 0.49 |
|  |  |  |  |  |  |
| --- | --- | --- | --- | --- | --- |
| **TABLE S8:** Mean relative abundance of the top four bacterial phyla on skin swabs taken from mothers from the four different tanks. Frogs in each tank are of similar age, as indicated by variable 'Mother Age'. | | | | | |

### **Table S9: ANCOMBC2 results**

| **Genus** | **Log fold change** | **Standard error** | **W statistic** | **p-value** | **q-value** |
| --- | --- | --- | --- | --- | --- |
| Rhizobium Genera | 5.64 | 0.65 | 8.64 | 5.81 × 10^−18^ | 4.41 × 10^−16^ |
| Bosea | 3.22 | 0.70 | 4.58 | 4.75 × 10^−6^ | 1.20 × 10^−4^ |
| Lachnospir. NK4A136 | 2.92 | 0.74 | 3.94 | 8.08 × 10^−5^ | 8.77 × 10^−4^ |
| Fluviicola | 2.73 | 0.87 | 3.14 | 1.71 × 10^−3^ | 8.12 × 10^−3^ |
| Delftia | 2.45 | 0.78 | 3.14 | 1.70 × 10^−3^ | 8.12 × 10^−3^ |
| Lactobacillus | 2.04 | 0.62 | 3.31 | 9.28 × 10^−4^ | 5.88 × 10^−3^ |
| Ralstonia | 1.74 | 0.48 | 3.60 | 3.14 × 10^−4^ | 2.65 × 10^−3^ |
| Dubosiella | 1.51 | 0.63 | 2.41 | 1.59 × 10^−2^ | 5.02 × 10^−2^ |
| Lachnoclostridium | 1.29 | 0.61 | 2.12 | 3.41 × 10^−2^ | 8.94 × 10^−2^ |
| Phyllobacterium | 1.23 | 0.54 | 2.29 | 2.23 × 10^−2^ | 6.13 × 10^−2^ |
| Staphylococcus | 1.18 | 0.58 | 2.04 | 4.15 × 10^−2^ | 1.05 × 10^−1^ |
| Klebsiella | 1.15 | 0.51 | 2.28 | 2.26 × 10^−2^ | 6.13 × 10^−2^ |
| [Eubacterium] xylanophilum group | 1.12 | 0.58 | 1.93 | 5.35 × 10^−2^ | 1.31 × 10^−1^ |
| Lachnospiraceae UCG-001 | 1.12 | 0.63 | 1.78 | 7.50 × 10^−2^ | 1.58 × 10^−1^ |
| Turicibacter | 1.03 | 0.59 | 1.75 | 8.05 × 10^−2^ | 1.61 × 10^−1^ |
| Marvinbryantia | 1.02 | 0.56 | 1.81 | 6.98 × 10^−2^ | 1.56 × 10^−1^ |
| Variovorax | 0.99 | 0.52 | 1.90 | 5.72 × 10^−2^ | 1.36 × 10^−1^ |
| Streptococcus | 0.91 | 0.50 | 1.79 | 7.30 × 10^−2^ | 1.58 × 10^−1^ |
| Prevotella\_7 | 0.88 | 0.50 | 1.76 | 7.82 × 10^−2^ | 1.61 × 10^−1^ |
| Achromobacter | 0.83 | 0.45 | 1.84 | 6.59 × 10^−2^ | 1.52 × 10^−1^ |
| Corynebacterium | 0.79 | 0.53 | 1.50 | 1.32 × 10^−1^ | 2.19 × 10^−1^ |
| Kocuria | 0.77 | 0.46 | 1.69 | 9.15 × 10^−2^ | 1.72 × 10^−1^ |
| Lawsonella | 0.72 | 0.47 | 1.53 | 1.26 × 10^−1^ | 2.17 × 10^−1^ |
| Aquabacterium | −0.66 | 0.43 | −1.52 | 1.30 × 10^−1^ | 2.19 × 10^−1^ |
| Deinococcus | −0.72 | 0.46 | −1.56 | 1.20 × 10^−1^ | 2.17 × 10^−1^ |
| Pseudoduganella | −0.73 | 0.47 | −1.54 | 1.23 × 10^−1^ | 2.17 × 10^−1^ |
| Terrisporobacter | −0.78 | 0.46 | −1.68 | 9.25 × 10^−2^ | 1.72 × 10^−1^ |
| Pedobacter | −1.05 | 0.61 | −1.72 | 8.51 × 10^−2^ | 1.66 × 10^−1^ |
| Clostridium s.s. 13 | −1.13 | 0.47 | −2.38 | 1.74 × 10^−2^ | 5.29 × 10^−2^ |
| Arcicella | −1.17 | 0.50 | −2.33 | 1.96 × 10^−2^ | 5.73 × 10^−2^ |
| Epulopiscium | −1.28 | 0.47 | −2.70 | 6.86 × 10^−3^ | 2.48 × 10^−2^ |
| Paracoccus | −1.47 | 0.53 | −2.76 | 5.79 × 10^−3^ | 2.20 × 10^−2^ |
| Alkanindiges | −1.49 | 0.51 | −2.91 | 3.57 × 10^−3^ | 1.51 × 10^−2^ |
| Laribacter | −1.52 | 0.45 | −3.36 | 7.87 × 10^−4^ | 5.44 × 10^−3^ |
| Clostridium s.s. 1 | −1.56 | 0.58 | −2.68 | 7.41 × 10^−3^ | 2.56 × 10^−2^ |
| Undibacterium | −1.61 | 0.66 | −2.44 | 1.46 × 10^−2^ | 4.82 × 10^−2^ |
| Paraclostridium | −1.62 | 0.55 | −2.94 | 3.26 × 10^−3^ | 1.46 × 10^−2^ |
| Massilia | −1.64 | 0.50 | −3.26 | 1.10 × 10^−3^ | 5.99 × 10^−3^ |
| Pseudomonas | −1.76 | 0.62 | −2.85 | 4.42 × 10^−3^ | 1.77 × 10^−2^ |
| Ideonella | −1.90 | 0.58 | −3.28 | 1.02 × 10^−3^ | 5.98 × 10^−3^ |
| Paucibacter | −2.27 | 0.54 | −4.24 | 2.25 × 10^−5^ | 3.42 × 10^−4^ |
| Flavobacterium | −2.51 | 0.68 | −3.67 | 2.46 × 10^−4^ | 2.34 × 10^−3^ |
| Rheinheimera | −2.54 | 0.58 | −4.40 | 1.10 × 10^−5^ | 2.08 × 10^−4^ |
| Vogesella | −2.64 | 0.75 | −3.53 | 4.23 × 10^−4^ | 3.21 × 10^−3^ |
| Romboutsia | −2.76 | 0.60 | −4.63 | 3.66 × 10^−6^ | 1.20 × 10^−4^ |
| Acinetobacter | −2.83 | 0.69 | −4.09 | 4.31 × 10^−5^ | 5.46 × 10^−4^ |
|  |  |  |  |  |  |
| --- | --- | --- | --- | --- | --- |
| **TABLE S9:** Significant results of ANCOMBC2 analysis to identify genera that are differentially abundant between mother skin swab and tadpole tail samples. Red text indicates a statistically significant p-value. Taxa in bold text are those found to be differentially abundant by both MaAsLin2 and ANCOMBC2. q-value = p-value adjusted with Benjamini-Hochberg correction. p < 0.25 | | | | | |

### **Table S10: MaAsLin2 results**

| **Genus** | **Coefficient** | **Standard error** | **N** | **N.not.0** | **p-value** | **q-value** |
| --- | --- | --- | --- | --- | --- | --- |
| Rhizobium Genera | 7.77 | 0.51 | 48 | 40 | 0.00 | 3.00 × 10^−17^ |
| Rheinheimera | −4.86 | 0.55 | 48 | 15 | 3.76 × 10^−11^ | 9.39 × 10^−10^ |
| Romboutsia | −4.44 | 0.61 | 48 | 18 | 3.84 × 10^−8^ | 5.03 × 10^−7^ |
| Laribacter | −1.92 | 0.29 | 48 | 10 | 4.02 × 10^−8^ | 5.03 × 10^−7^ |
| Acinetobacter | −4.96 | 0.70 | 48 | 31 | 5.33 × 10^−8^ | 5.33 × 10^−7^ |
| Epulopiscium | −3.30 | 0.52 | 48 | 10 | 1.00 × 10^−7^ | 8.35 × 10^−7^ |
| Paucibacter | −2.04 | 0.30 | 48 | 11 | 1.36 × 10^−7^ | 9.72 × 10^−7^ |
| Flavobacterium | −4.23 | 0.68 | 48 | 33 | 5.93 × 10^−7^ | 3.71 × 10^−6^ |
| Bosea | 4.83 | 0.83 | 48 | 37 | 6.68 × 10^−7^ | 3.71 × 10^−6^ |
| Clostridium s.s. 13 | −2.57 | 0.46 | 48 | 9 | 1.26 × 10^−6^ | 5.34 × 10^−6^ |
| Vogesella | −4.42 | 0.73 | 48 | 24 | 1.10 × 10^−6^ | 5.34 × 10^−6^ |
| Massilia | −3.34 | 0.56 | 48 | 13 | 1.28 × 10^−6^ | 5.34 × 10^−6^ |
| Arcicella | −2.88 | 0.50 | 48 | 10 | 2.54 × 10^−6^ | 9.75 × 10^−6^ |
| Paracoccus | −2.73 | 0.51 | 48 | 11 | 3.07 × 10^−6^ | 1.10 × 10^−5^ |
| Pseudomonas | −3.99 | 0.71 | 48 | 39 | 3.52 × 10^−6^ | 1.17 × 10^−5^ |
| Cellulosilyticum | −2.48 | 0.47 | 48 | 8 | 3.78 × 10^−6^ | 1.18 × 10^−5^ |
| Deinococcus | −2.18 | 0.42 | 48 | 9 | 4.66 × 10^−6^ | 1.37 × 10^−5^ |
| Ideonella | −2.32 | 0.45 | 48 | 9 | 5.89 × 10^−6^ | 1.63 × 10^−5^ |
| dgA-11 gut group | −2.15 | 0.43 | 48 | 10 | 9.03 × 10^−6^ | 2.38 × 10^−5^ |
| Paraclostridium | −1.70 | 0.34 | 48 | 8 | 1.23 × 10^−5^ | 3.06 × 10^−5^ |
| Aquabacterium | −1.37 | 0.28 | 48 | 8 | 1.47 × 10^−5^ | 3.49 × 10^−5^ |
| Clostridium s.s. 1 | −4.17 | 0.82 | 48 | 23 | 1.80 × 10^−5^ | 4.09 × 10^−5^ |
| Alkanindiges | −1.99 | 0.42 | 48 | 11 | 2.20 × 10^−5^ | 4.78 × 10^−5^ |
| Lachnospir. NK4A136 | 3.27 | 0.76 | 48 | 26 | 9.07 × 10^−5^ | 1.89 × 10^−4^ |
| Ralstonia | 1.29 | 0.29 | 48 | 18 | 9.75 × 10^−5^ | 1.95 × 10^−4^ |
| Halomonas | −1.45 | 0.36 | 48 | 8 | 2.11 × 10^−4^ | 4.06 × 10^−4^ |
| Desulfovibrio | −2.06 | 0.55 | 48 | 9 | 4.72 × 10^−4^ | 8.74 × 10^−4^ |
| Psychrobacter | −1.38 | 0.37 | 48 | 10 | 5.22 × 10^−4^ | 9.33 × 10^−4^ |
| Enhydrobacter | −2.34 | 0.61 | 48 | 11 | 5.63 × 10^−4^ | 9.70 × 10^−4^ |
| Lactobacillus | 2.62 | 0.71 | 48 | 22 | 6.67 × 10^−4^ | 1.11 × 10^−3^ |
| Delftia | 3.60 | 0.96 | 48 | 26 | 7.37 × 10^−4^ | 1.19 × 10^−3^ |
| Fluviicola | 4.45 | 1.31 | 48 | 22 | 1.91 × 10^−3^ | 2.98 × 10^−3^ |
| Undibacterium | −2.04 | 0.62 | 48 | 19 | 2.00 × 10^−3^ | 3.04 × 10^−3^ |
| Sphingomonas | −2.14 | 0.68 | 48 | 30 | 2.80 × 10^−3^ | 4.12 × 10^−3^ |
| Alistipes | −2.03 | 0.66 | 48 | 14 | 4.23 × 10^−3^ | 6.05 × 10^−3^ |
| Brevundimonas | −1.01 | 0.37 | 48 | 8 | 1.09 × 10^−2^ | 1.51 × 10^−2^ |
| Pedobacter | −1.54 | 0.59 | 48 | 25 | 1.20 × 10^−2^ | 1.63 × 10^−2^ |
| Phyllobacterium | 0.83 | 0.37 | 48 | 9 | 3.15 × 10^−2^ | 4.14 × 10^−2^ |
| Dubosiella | 0.90 | 0.40 | 48 | 9 | 3.42 × 10^−2^ | 4.38 × 10^−2^ |
| Klebsiella | 0.59 | 0.31 | 48 | 9 | 6.68 × 10^−2^ | 8.35 × 10^−2^ |
| Staphylococcus | 1.20 | 0.69 | 48 | 15 | 9.35 × 10^−2^ | 1.14 × 10^−1^ |
| Achromobacter | 0.38 | 0.23 | 48 | 8 | 1.09 × 10^−1^ | 1.30 × 10^−1^ |
| Turicibacter | 0.96 | 0.59 | 48 | 24 | 1.15 × 10^−1^ | 1.34 × 10^−1^ |
|  |  |  |  |  |  |  |
| --- | --- | --- | --- | --- | --- | --- |
| **TABLE S10:** Significant results of MaAsLin2 analysis to identify genera that are differentially abundant between mother skin swab and tadpole tail samples. coef = model coefficient (effect size). stderr = coefficient standard error. Red text indicates a statistically significant p-value. Taxa in bold text are those found to be differentially abundant by both MaAsLin2 and ANCOMBC2. N = number of samples. N.not.0 = number of samples where the taxon count was non-zero. q-value = p-value adjusted with Benjamini-Hochberg correction. p < 0.25 | | | | | | |

### Table S11: FEAST results summary

| **Source** | **Max** | **Mean** | **Min** | **SD** |
| --- | --- | --- | --- | --- |
| Mother skin swab | | | | |
| --- | --- | --- | --- | --- |
| Tank swab | 0.01 | 0.00 | 0.00 | 0.00 |
| Tank water | 0.98 | 0.61 | 0.16 | 0.29 |
| Unknown | 0.84 | 0.39 | 0.02 | 0.29 |
| Egg jelly wash | | | | |
| Mother skin swab | 0.99 | 0.73 | 0.39 | 0.20 |
| Unknown | 0.61 | 0.27 | 0.01 | 0.20 |
| Tadpole tail | | | | |
| Egg jelly wash | 0.99 | 0.60 | 0.12 | 0.26 |
| Unknown | 0.88 | 0.40 | 0.01 | 0.26 |
| Tadpole plate media | | | | |
| Egg jelly wash | 0.16 | 0.02 | 0.00 | 0.03 |
| Tadpole tail | 0.99 | 0.68 | 0.00 | 0.32 |
| Unknown | 0.99 | 0.31 | 0.00 | 0.31 |
|  |  |  |  |  |
| --- | --- | --- | --- | --- |
| **TABLE S11:** Summary of results of FEAST microbial source tracking analysis to track the flow of microbial communities through the Xenopus life cycle and between frogs and the environment. Results are divided into groups comprised of a sink (i.e. sample type receiving microbiota - shown in bold) and the potential source samples for that sink. Numbers provided are proportions; Max = maximum values, Min = minimum value, SD = standard deviation. | | | | |
